# Where the elements go when it rains: carbon and nutrient stocks of soil amended with stabilised organic matter

**DOI:** 10.1101/015453

**Authors:** E-L Ng, C.R. Schefe, T.R. Cavagnaro, A.F Patti

## Abstract

The addition of compost to the soil can increase C and nutrient levels, but how these are affected by different rainfall regimes remain unknown. This study determines the carbon (C), nitrogen (N) and phosphorus (P) stocks after drying and rewetting events in a compost-amended grassland ecosystem at 3 depths (0-5, 5-10, 10-30cm). Compost addition consistently increased C, N and P stocks despite altered rainfall but these improvements can only be detected by careful consideration of soil depths.

## Introduction

Grasslands cover approximately 40% of terrestrial surface and store a third of terrestrial carbon, of which considerably more is stored in soils than vegetation (*White, Murray, and Rohweder* 2000). However, many of the worlds grasslands are degraded and organic input has been proposed as a means of replenishing soil organic matter in grasslands (*Jackson et al*. 2004; *Asner et al*. 2004; *Powlson et al*. 2012). As it stands, the fate of organic matter added to grasslands remains unclear (*Ryals et al*. 2014). The decomposition of organic matter in soils is regulated by soil water and nutrient availability, temperature and pH (*Clark et al*. 2009; *Borken and Matzner* 2009; *Elhottová et al*. 2006). With much of the world projected to experience changes in rainfall regimes under climate change (*IPCC* 2013), there may be consequences for decomposition and elemental cycling. The few studies that examined the response of soil processes to simultaneous drivers of global change have found surprising antagonistic interactions led to effects much less severe than expected (*Dieleman et al*. 2012; *Eisenhauer et al*. 2012; *Dukes et al*. 2005). Here, we address this gap by quantifying soil C, N and P stocks after drying and rewetting events in a compost amended grassland soil from southeastern Australia.

## Methods

Intact soil cores (15cm diameter×40cm deep) were collected from an intensively grazed grassland (Lat-38.010551/Long-145.472542). Details of the site, rain treatments and terrestrial model ecosystem setup are provided in Ng *et al*. (in review). In short, compost was applied onto the surface of each soil core at 30 t/ha (dry mass). Based on long-term weather records, each core under normal rain received 47.8mm, 65.0mm and 83.2mm of simulated rainfall; each core under drying received 4.0mm, 18.4mm and 12.4mm for March, April and May respectively; the rewetting treatment was the same as the drying treatment, but it also received an additional 150mm three days before experiment ends. The cores were organised into randomised complete block design, with each treatment appearing once in each of the 5 blocks. The cores were destructively sampled at 0-5, 5-10 and 10-30cm, 3 months after compost addition. Samples were sieved to 0.2cm, air-dried and ground for elemental analysis. Total C, N and P were measured on all soil layers (see Ng et al. in review).

Data were analysed using randomised complete block design ANOVA. Where assumptions of normality and homoscedasticity are not met, transformations are carried out. These results were compared to results of untransformed data. As they are the same, results of the untransformed data are shown. Post-hoc multiple comparisons were carried out using least significant difference (LSD) test with *p*-values adjusted using bonferroni. Data analysis was carried out on R 2.15.1 (*R Core Team* 2012) using agricolae package (*Mendiburu* 2012) for LSD test.

## Results and discussion

Carbon stocks were altered with compost amendment only at 5 to 10cm (Fig.1; F_1,22_=6.7, *p*=0.017). Specifically, C stocks were higher in amended soils (mean±standard error=2.8±0.1kg/m^2^) compared to unamended soils (2.5±0.2kg/m^2^) in the 5 to 10cm soil layer, but not the other soil layers. Rainfall treatment did not have any significant effects on the C stocks nor did it alter the effect of compost on C stock at all depths. Cumulative C stock from 0-30cm was similar across compost and rain treatments.

**Fig. 1.**
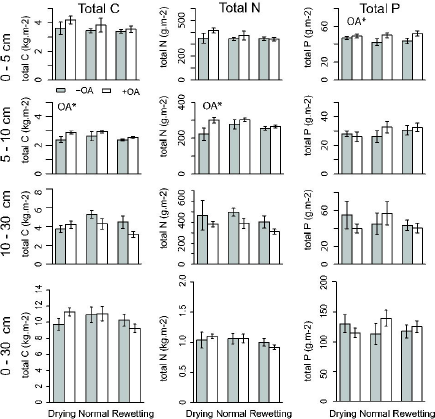
Effects of OA on total C, N and P at 0-5 cm, 5-10 cm, 10-30 cm and 0-30 cm under drying, normal and rewetting treatments. Significant levels are \**p* < 0.05, ** *p* < 0.01, *** *p* < 0.001 (refer in-text for test statistics).

Nitrogen stock was also altered with compost amendment at 5 to 10cm (Fig.1; F_1,22_=5.4, *p*=0.030). Soil N stocks were higher in amended soils compared to unamended soils. Rainfall treatment did not have any effects on the N stocks nor did it alter the effect of compost on N stock at all depths. Cumulative N stock (0-30cm) was similar across compost and rain treatments.

Soil P stocks were altered with compost amendment at only 0 to 5cm (Fig.1; F_1,22_=6.1, *p*=0.021), where compost amended soils had higher total P. Rainfall treatment did not have any effects on the P stocks nor did it alter the effect of compost on P stock at all depths. Cumulative P stock from 0 to 30cm was similar across OA and rain treatments.

A single application of organic amendment can improve grassland soil organic matter for up to 14 years after application (*Ippolito et al*. 2010; *Ryals et al*. 2014). However, as rainfall affects soil physicochemical properties such as soil water dynamics and the movement of soil organic matter along the profile, and these changes in turn affects the soil biota and their feedback to the elemental cycling (*Knapp et al*. 2008; *Borken and Matzner* 2009; *Bell, Sherry, and Luo* 2010; *Berhe et al*. 2012), altered rainfall may modify the elemental cycling in the ecosystem. In our grassland with sandy soil, all the added C, N and P have been retained in the 0 to 30cm during this 3 month experiment (data not shown). It will be important to assess the longer term fate of compost-derived C, N and P in these soils under different rainfall treatments.

Although the total stocks did not change with rainfall, it is important to note that available nutrients were different across the rain treatments; we observed higher NH_4_^+^ in drying and rewetting treatment compared to normal rain and NO_3_^-^ was highest in the rewetting treatment, followed by the drying and normal rain treatments at 0 to 5cm (see Ng et al. in review). But similarly to results of the total stocks, there was no interaction effect between the rainfall and compost treatments on mineral N. The P accumulation at the 0 to 5cm with addition of compost indicated low P availability, likely due to high contents of calcium, aluminium and iron in this soil (data not shown). Given primary productivity depends on available nutrients, changes in this pool with rainfall is important.

When C, N and P stocks were pooled over all depths no differences among treatments were observed. This highlights the importance of taking sampling depth into consideration when assessing changes in C stocks. A similar response has been observed when tillage effects on soil C have been studied, with differences detectable at 0-30cm, but not 0-15cm, while the effects of perennial forage were detectable at all sampling depths to 60cm (*VandenBygaart et al*. 2011). These changes in elemental distribution across the soil profile may have important implications on the outcome of competition between plants, and between plants and soil fauna that utilise those resources, and degree of nutrient leaching (*Franzluebbers and Hons* 1996).

Green waste-derived compost provides a genuine possibility to increase soil C while diverting biomass from landfill (*Powlson et al*. 2012). Based on our findings, the elements (C, N and P) of soil amended with stabilised organic matter (compost) stayed in the sandy grassland even when the rain poured down.

## Acknowledgements

This work was supported by funding from the Monash Sustainability Institute Australia, Brown Coal Innovation Australia and the Australian Research Council (ARC). NEL wishes to thank Pablo Galaviz, Mattia Pierangelini, Alicia Brown, Mani Shresta for assisting field and lab work. TRC also thanks the ARC for the award of a Future Fellowship (FT120100463).

## References

Asner, G. P., A. J. Elmore, L. P. Olander, R. E. Martin, and T. Harris (2004): Grazing systems, ecosystem responses, and global change. Annu. Rev. Env. Resour. 29, 261.

Bell, Jesse E., Rebecca Sherry, and Luo Yiqi (2010): Changes in soil water dynamics due to variation in precipitation and temperature: An ecohydrological analysis in a tallgrass prairie. Water Resour. Res. 46, W03523.

Berhe, Asmeret, K. Suttle, Sarah Burton, and Jillian Banfield (2012): Contingency in the direction and mechanics of soil organic matter responses to increased rainfall. Plant Soil 358, 371.

Borken, W., and E. Matzner (2009): Reappraisal of drying and wetting effects on C and N mineralization and fluxes in soils. Glob. Change Biol. 15, 808.

Clark, J. S., J. H. Campbell, H. Grizzle, V. Acosta-Martìnez, and J. C. Zak (2009): Soil microbial community response to drought and precipitation variability in the chihuahuan desert. Microb. Ecol. 57, 248.

Dieleman, W. I. J., S. Vicca, F. A. Dijkstra, F. Hagedorn, M. J. Hovenden, K. S. Larsen, J. A. Morgan, A. Volder, C. Beier, J. S. Dukes, J. King, S. Leuzinger, S. Linder, Y. Luo, R. Oren, P. De Angelis, D. Tingey, M. R. Hoosbeek, and I. A. Janssens (2012): Simple additive effects are rare: A quantitative review of plant biomass and soil process responses to combined manipulations of CO 2 and temperature. Glob. Change Biol. 18, 2681.

Dukes, J. S., N. R. Chiariello, E. E. Cleland, L. A. Moore, M. Rebecca Shaw, S. Thayer, T. Tobeck, H. A. Mooney, and C. B. Field (2005): Responses of grassland production to single and multiple global environmental changes. PLoS Biol. 3, pe319.

Eisenhauer, Nico, Simone Cesarz, Robert Koller, Kally Worm, and Peter B. Reich (2012): Global change below ground: impacts of elevated CO2, nitrogen and summer drought on soil food webs and biodiversity. Glob. Change Biol. 18, 435.

D. Elhottová, V. Krištůfek, J. Tříska, V. Chrastný, E. Uhlířová, J. Kalčík, and T. Picek (2006): Immediate impact of the flood (Bohemia, August 2002) on selected soil characteristics. Water. Air. Soil Pollut. 173, 177.

Franzluebbers, A. J., and F. M. Hons (1996): Soil-profile distribution of primary and secondary plant-available nutrients under conventional and no tillage. Soil Tillage Res. 39, 229.

IPCC (2013): Climate Change 2013: The Physical Science Basis. Contribution of Working Group I to the Fifth Assessment Report of the Intergovernmental Panel on Climate Change. In T. F. Stocker, D. Qin, G.-K. Plattner, M. Tignor, S. K. Allen, J. Boschung, A. Nauels, Y. Xia, V. Bex, and P. M. Midgley, 1535.

Ippolito, J. A., K. A. Barbarick, M. W. Paschke, and R. B. Brobst (2010): Infrequent composted biosolids applications affect semi-arid grassland soils and vegetation. J, Environ. Manage. 91, 1123.

Jackson, L. E., I. Ramirez, R. Yokota, S. A. Fennimore, S. T. Koike, D. M. Henderson, W. E. Chaney, F. J. Calderόn, and K. Klonsky (2004): On-farm assessment of organic matter and tillage management on vegetable yield, soil, weeds, pests, and economics in California. Agr. Ecosyst. Environ. 103, 443.

Knapp, A. K., C. Beier, D. D. Briske, A. T. Classen, Y. Luo, M. Reichstein, M. D. Smith, S. D. Smith, J. E. Bell, P. A. Fay, J. L. Heisler, S. W. Leavitt, R. Sherry, B. Smith, and E. Weng (2008): Consequences of More Extreme Precipitation Regimes for Terrestrial Ecosystems. Bioscience 58, 811.

Mendiburu, Felipe (2012): Agricolae: Statistical Procedures for Agricultural Research,

Powlson, D. S., A. Bhogal, B. J. Chambers, K. Coleman, A. J. Macdonald, K. W. T. Goulding, and A. P. Whitmore (2012): The potential to increase soil carbon stocks through reduced tillage or organic material additions in England and Wales: A case study. Agr. Ecosyst. Environ. 146, 23.

R Core Team (2012): R: A language and environment for statistical computing. R Foundation for Statistical Computing: Vienna, Austria,

Ryals, R., M. Kaiser, M. S. Torn, A. A. Berhe, and W. L. Silver (2014): Impacts of organic matter amendments on carbon and nitrogen dynamics in grassland soils. Soil Biol. Biochem. 68, 52.

VandenBygaart, A. J., E. Bremer, B. G. McConkey, B. H. Ellert, H. H. Janzen, D. A. Angers, M. R. Carter, C. F. Drury, G. P. Lafond, and R. H. McKenzie (2011): Impact of sampling depth on differences in soil carbon stocks in long-term agroecosystem experiments. Soil Sci. Soc. Am. J. 75, 226.

White, R., S. Murray, and M. Rohweder (2000): Pilot Analysis of Global Ecosystems: Grassland Ecosystems. World Resources Institute: Washington D.C., USA, 69.

